# The entangled world of DNA quadruplex folds

**DOI:** 10.1101/2024.04.17.589856

**Authors:** Sruthi Sundaresan, Patil Pranita Uttamrao, Purnima Kovuri, Thenmalarchelvi Rathinavelan

## Abstract

DNA quadruplexes take part in many biological functions. It takes up a variety of folds based on the sequence and environment. Here, a meticulous analysis of experimentally determined 392 quadruplex structures (388 PDB IDs) deposited in PDB is carried out. The analysis reveals the modular representation of the quadruplex folds. 48 unique quadruplex motifs (whose diversity arises out of the propeller, bulge, diagonal, and lateral loops that connect the quartets) are identified, leading to simple to complex inter-/intra-molecular quadruplex folds. These structural two-layered motifs are further classified into 33 continuous and 15 discontinuous motifs. The discontinuous motifs cannot further be classified into parallel, antiparallel, or hybrid as one or more guanines of the adjacent quartets are not connected. While the continuous motifs can be extended to a quadruplex fold, the discontinuous motif requires additional loop(s) to complete a fold, as illustrated here with examples. Similarly, the higher-order quadruplex folds can also be represented by continuous or discontinuous motifs or their combinations. Such a modular representation of the quadruplex folds may assist in custom engineering of quadruplexes, designing motif-based drugs, and the prediction of quadruplex structure. Further, it could facilitate understanding the role of quadruplexes in biological functions and diseases.

## Introduction

Besides the right-handed double helical structure [1, 2], nucleic acids can fold into a variety of secondary structures like hairpin, triplex, quadruplex, i-motif, *etc* [3–9]. Multiple pieces of evidence unfold the instrumental role of quadruplex in regulating the biological processes across all the domains of life [10–22]. This class of thermodynamically stable alternative structure encompasses four guanine-rich strands [23] (thus the name quadruplex or tetraplex), stabilized by planar G-tetrads (**Figure 1A**) that stack on each other (**Figure 1B, 1C**). The inward orientation of the carbonyl groups of the guanines in the G-quartet creates an electronegative repulsion. Thus, there is a requirement for mono- or di-valent cations to stabilize the G-quartets [24, 25]. Although the quartet structure was originally found to be formed only by four guanines, recent evidence shows the possibility of non-G-quartets in the middle of G-quartets [26, 27].

**Figure 1.**
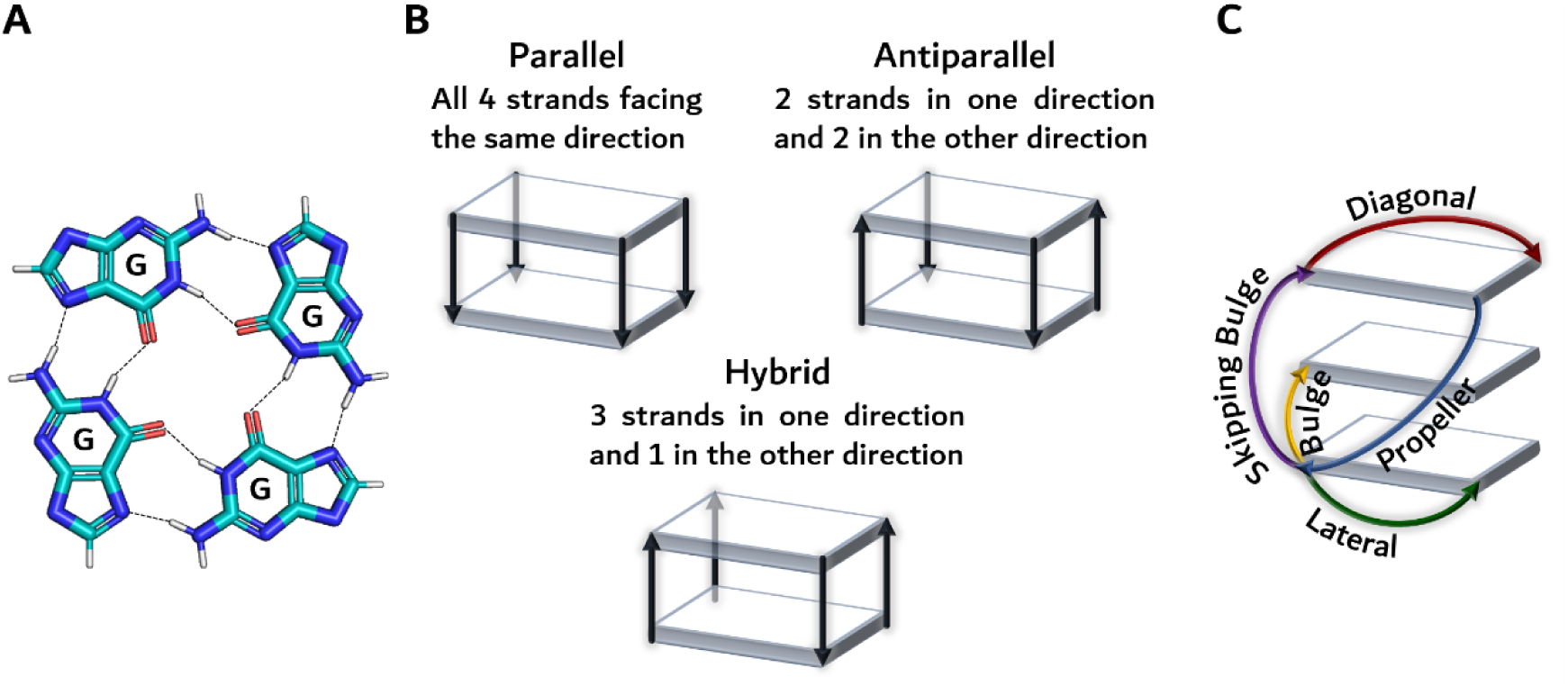
Nomenclature used in this study for describing the quadruplex folds. A) Hydrogen bonding (dotted line) pattern of a G-quartet, B) Schematic illustration of parallel, antiparallel, hybrid quadruplex folds, C) Loops observed in quadruplex folds. See text for details.

Sequences rich in Gs (which are prone to form thermodynamically favorable quadruplex structures) are found to be pervasive in the human genome [18, 19, 28, 29] and have important regulatory roles that are influenced by their genomic location [11]. For instance, sequences prone to form quadruplexes are enriched (nearly 50%) in the human gene promoters [29], suggestive of their role in the genetic transactions [23, 30], and also play a vital role in governing gene expression and genomic stability [18, 19, 28, 29]. Notably, the quadruplex structures formed in the promoter regions of oncogenes like KRAS [31], c-Myc [32], *etc.,* negatively regulate them. These structures are also present in the telomeres of eukaryotes [33] and are shown to inhibit telomere extension [34]. Quadruplex structures have been identified in the recombination hotspots of the human genome, insisting on their role in recombination [35]. The prevalence of quadruplex-forming sequences in the human transcriptome further facilitates the formation of RNA quadruplex structures [27]. RNA quadruplexes take part not only in mRNA cap-dependent [36] and independent [37] translation but also in alternative splicing [38], mitochondrial transcription termination [39], mRNA localization [40], and maturation of miRNAs [17, 41]. Besides independently forming quadruplex structures, RNA and DNA can together form an RNA-DNA hybrid quadruplex. Such a hybrid quadruplex structure formed between the template DNA and nascent mRNA strand is shown to play a role in transcription termination [37]. Recently, there have been a number of convincing studies on the role of quadruplex structures in misregulating the RNA biology of repeat expansion-associated neurodegenerative disorders and causing neuroglial pathogenies through non-conventional repeat-associated non-AUG (RAN) translation [42–45]. G-rich sequences that are prone to form quadruplex structures are also found in the regulatory regions of microbes such as bacteria, fungi, and viruses. For example, a number of putative quadruplex-forming sequences are found in *Plasmodium falciparum* [10]. Quadruplex structures have also been shown to have a role in viral replication, modulating translation, and facilitating immune evasion [11]. Additionally, they play a role in modulating microbial transcription through radioresistance, antigenic variation, recombination, and latency [11].

Due to its participation in the aforementioned diverse biological functions, quadruplex structures act as disease biomarkers [46] and potential therapeutic targets: anticancer [34, 47], antimicrobial [11, 48], anticoagulant [49], and anti-neurodegenerative [47, 50]. Besides, G-quadruplex-forming aptamers play a role in drug delivery [51] and diagnoses [52]. There are several proven examples of the role of quadruplexes in nanotechnological applications, such as biosensors [53, 54], nanomotors [55], nanomaterials [56], origami scaffolds (to capture direct interaction between quadruplexes and proteins) [57], and nanowires in nanoelectronics [58].

Since quadruplex structures are intrinsically polymorphic in nature, they are greatly influenced by the sequence and environmental conditions like pH, cations, *etc*. Several research groups around the world are working on getting the atomistic details of these structures using X-ray crystallography, NMR, and cryo-EM [59–61]. However, only a few studies have summarized in detail the atomistic structures of quadruplexes [62–65]. Recently, this lab has reported ten unique folding topologies of RNA quadruplexes [27] after analyzing in detail the structures deposited in the protein databank (PDB) [60]. Considering the diverse therapeutic potential and nanotechnology applications of DNA quadruplexes, a systematic analysis of DNA quadruplex structures deposited in PDB has been carried out here. The information provided here may enlighten the topological diversity of DNA quadruplex folds in understanding not only their biological significance and therapeutics but also in the custom design of DNA quadruplex scaffolds for nanotechnological applications.

## Materials and methods

### DNA quadruplex structures

In order to understand the architecture of various DNA quadruplex folds, the cartesian coordinates of quadruplex structures were downloaded from PDB [60], NDB [61, 66], and ONQUADRO [59] using the keyword search "quadruplex" or "tetraplex" or "tetrad" or "quartet" or "G4-helices". After removing the redundant structures, 392 DNA quadruplexes (388 PDB IDs) were finally considered for analysis. Note that the intercalating quadruplex structures (PDB IDs: 1V3P, 2DZ7) (*viz.*, octaplex) are not considered for the analysis as it is beyond the scope of this article.

### Deriving structural two-layered motifs from the DNA quadruplex structures

Since the objective of this study is to show that the quadruplex structures or folds are modular in nature in such a way that can be represented in terms of small repetitive 3D structural units, a thorough analysis of 392 quadruplex PDB structures was carried out in this context. The 3D structural units, henceforth referred to as quadruplex motifs, were derived from the quadruplex folds by breaking the folds into two-layered structural blocks comprising two quartets.

### Conformational angle preference

The sugar-phosphate backbone and glycosyl conformational angles [α (C3’-P-O5’-C5’), β (P-O5’-C5’-C4’), γ (O5’-C5’-C4’-C3’), ε (C4’-O3’-C3’-P), ζ (O3’-C3’-P-O5’), χ (O4’-C1’-N9/N1-C6/C4)] of each quartet-forming residues in the quadruplex folds (PDBs) was taken from NDB. The conformational angle preference is reported based on the frequency of occurrence, which is individually normalized with respect to each fold, *viz.*, separately for antiparallel, parallel and hybrid which was then plotted as bar plot using Excel.

### Plots

The contour density plots that show the conformational angle preferences of the quartet and loop residues were generated with the help of Gnuplot [67].

## Results and Discussions

### General description of DNA quadruplex architecture

The nomenclatures used in the description of quadruplex folds are shown in **Figure 1**. **Figure 1B** represents the well-known three broad classifications of quadruplex based on the relative orientations of their four strands: parallel, antiparallel, and hybrid folds. In a parallel quadruplex fold, the directions of all four strands are the same. In the antiparallel fold, two strands are in one direction, and the other two strands are in the opposite direction. In contrast, a hybrid quadruplex fold has three strands in one direction and the other strand in the opposite direction. However, this nomenclature is insufficient to understand the quadruplex folds, as the internal loops can further complicate the quadruplex folds. There are four loops observed in the quadruplex folds: diagonal, lateral, propeller, and bulge **(Figure 1C)** [68]. The diagonal loop connects the guanines of the alternate strands in the terminal quartet, and the lateral loop connects the adjacent strand guanines of the terminal (same) quartet. The former leads to a change in the direction of the alternate strands, and the latter results in a change in the direction of the adjacent strands. Unlike the lateral and diagonal loops, the propeller and bulge loops connect two different quartets. In most situations, the propeller loop connects two terminal quartets that are located at the opposite ends of a quadruplex and connect the adjacent strands. Exceptionally, a propeller loop can also connect a terminal quartet to a non-terminal quartet involving the adjacent strands. The propeller loop connects the two adjacent strands in a parallel fashion. Unlike these three loops, the bulge loop connects the same strand, wherein it can be present between 2 adjacent (bulge, **Figure 1C**) or non-adjacent (skipping bulge, **Figure 1C**) quartets without and with changing the direction of the strand, respectively.

### Classification of quadruplex motifs

223 unique sequences from 392 DNA quadruplex structures (388 PDB IDs) reveal that there are 49 unique folds. Intriguingly, an in-depth analysis indicates that these quadruplex folds are modular in nature as they can be represented in terms of small repetitive 3D structural units, henceforth referred to as quadruplex motifs. These motifs can be classified into two broad categories (**Figure 2**): continuous and discontinuous. In the former, all four residues of each adjacent quartet are connected continuously by the phosphodiester bonds of the backbone with or without the bulge loops. In the latter, at least one of the four residues of each adjacent quartet is not connected by the phosphodiester bond. These two categories can further be classified into four classes (**Figures 3, 4**): monomer (made with a single strand) (**Figure 3A, 4A**), dimer (made up of 2 different strands) (**Figure 3B, 4B**), trimer (made up of 3 different strands) (**Figure 3C, 4C**) and tetramer (made up of 4 different strands) (**Figure 3D**). The tetramer motifs are found only under the continuous category. These classes of the continuous category can further be classified into parallel, antiparallel, and hybrid motif types based on the strand orientation (Refer to **Figure 1B**). There are 14, 9, 6, and 4 monomers (**Figure 3A**), dimers (**Figure 3B**), trimers (**Figure 3C**), and tetramers (**Figure 3D**) motifs, respectively, seen under the continuous motif category (**Figure 3**). Similarly, there are 6, 3, and 6 monomers (**Figure 4A**), dimers (**Figure 4B**), and trimers (**Figure 4C**) motifs, respectively, seen under the discontinuous motif category (**Figure 4**). Note that the discontinuous motifs cannot further be classified into parallel, antiparallel, or hybrid as one or more guanines of the adjacent quartets are not connected. Hereinafter, for the sake of description purpose, each motif is given a five-letter alphanumeric name as described below.

**Figure 2.**
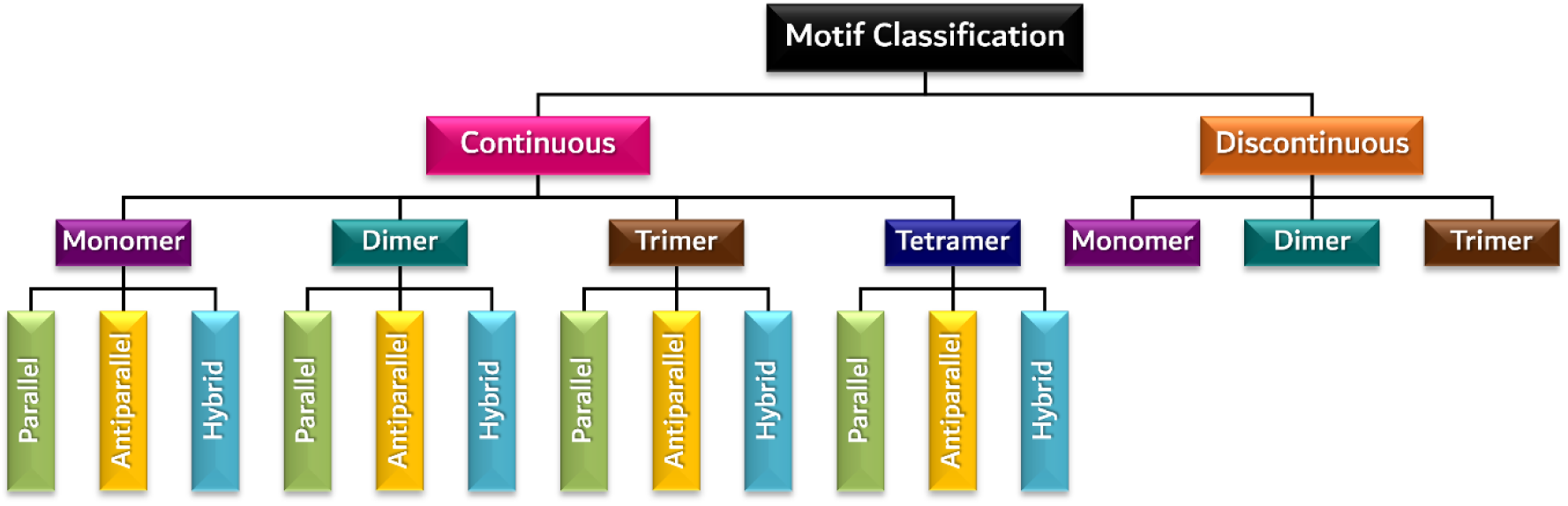
Quadruplex motif classification.

**Figure 3.**
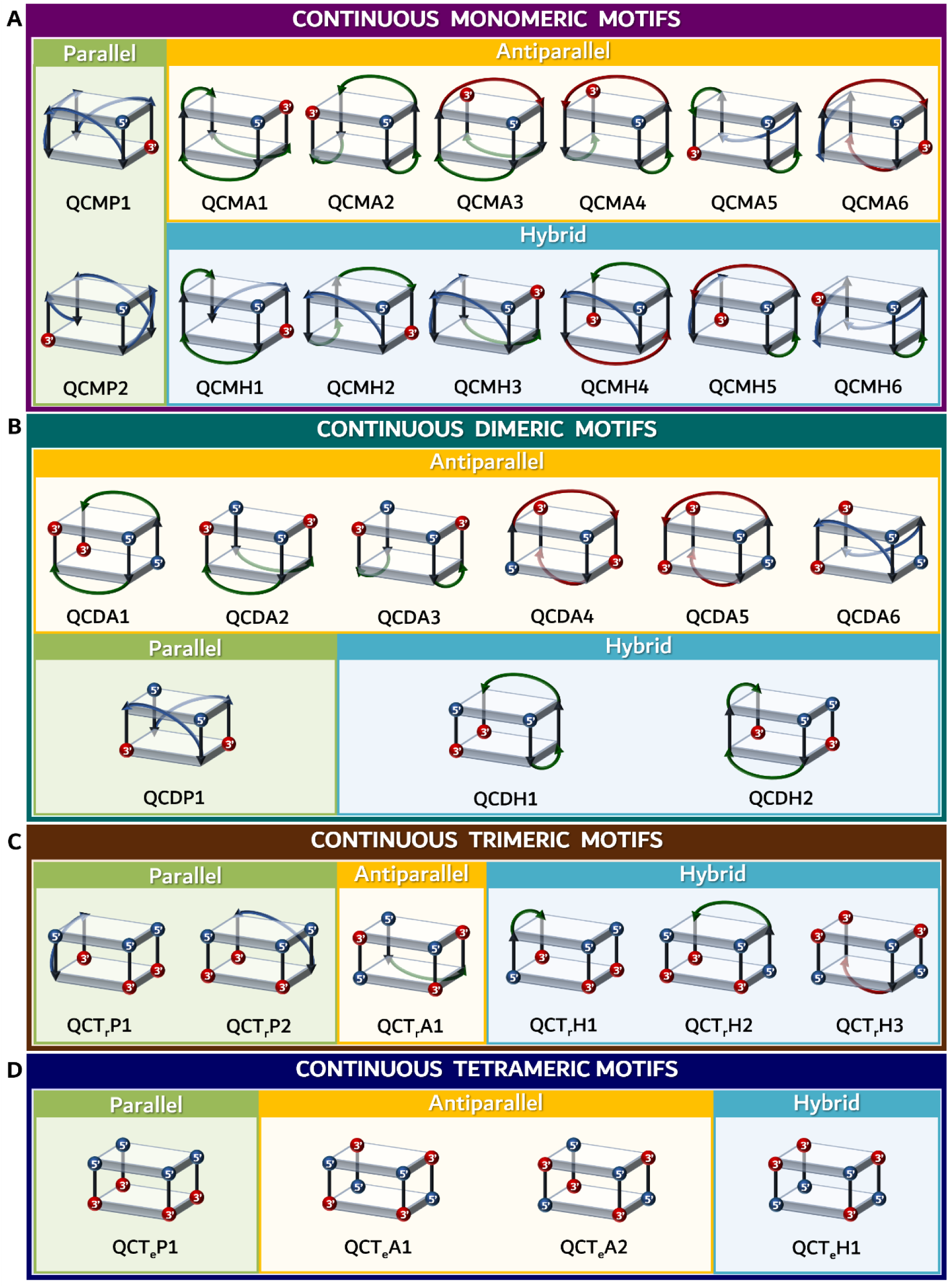
Schematic representation of the continuous quadruplex motifs. Note that the bulge loop is ignored in the representation as it wouldn’t affect the strand direction. Thus, for the representation purpose, the bulge loop residues are considered to be zero. Note that to maintain uniformity, always the 5’ end (blue-colored sphere) is kept at the top right and front face of the quadruplex motif. In the multimeric dimer, trimer, and tetramer, the 5’ end of any one of the strands is kept in the top right and front face of the motif. The red-colored sphere represents the 3’ end.

**Figure 4.**
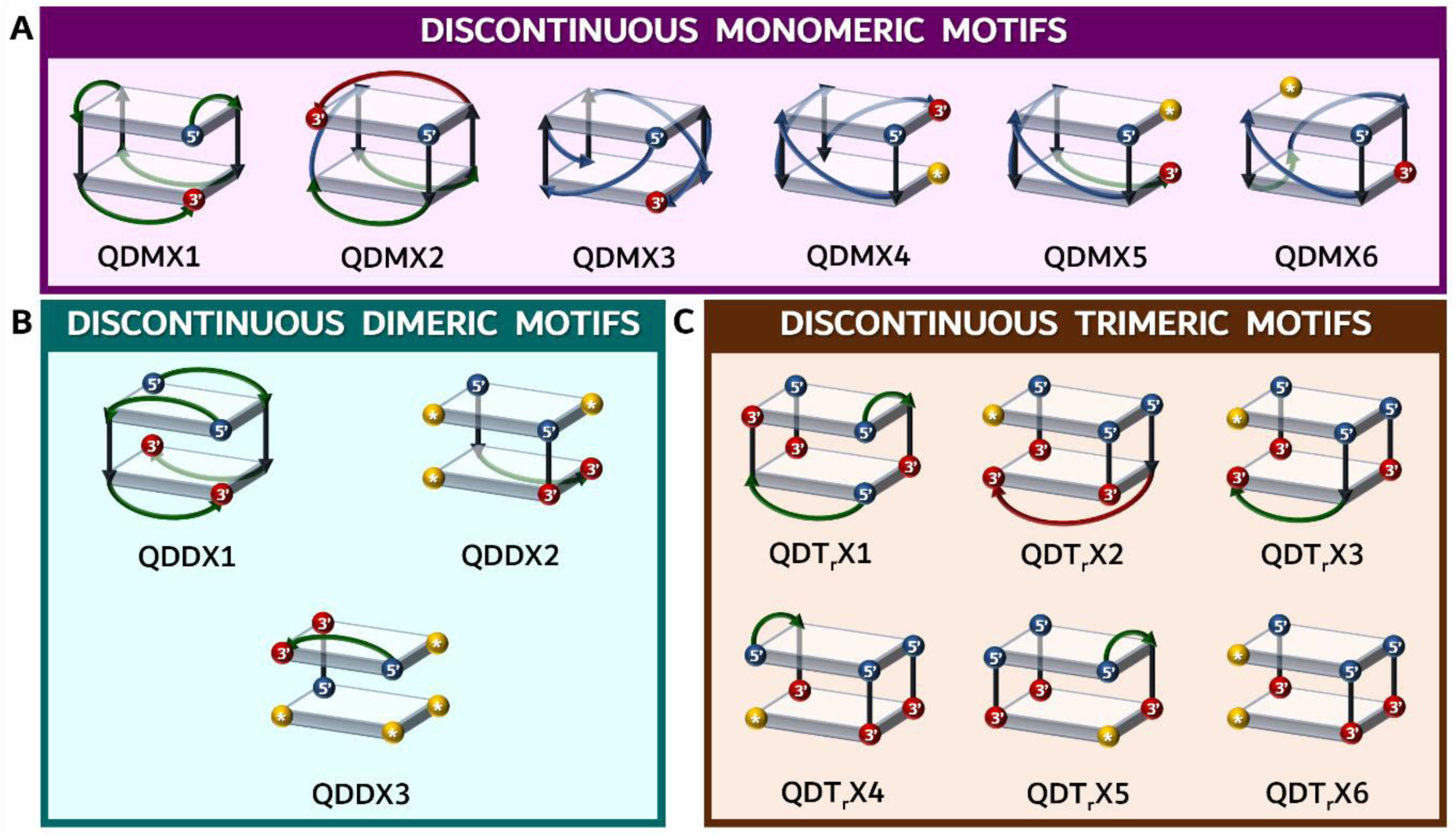
Schematic representation of the discontinuous quadruplex motifs. Note that the star enclosed by a golden color sphere represents the nucleotide monomer whose orientation can be of any direction (*viz.*, 5’ or 3’). Such single nucleotides are not considered independent strands, thus, not used in defining the quadruplex motif multimer.

The first letter starts with the alphabet "Q", representing the quadruplex (Q) followed by the category (C (continuous) or D (discontinuous)) followed by the class (M (monomer) or D (dimer) or T_r_ (trimer) or T_e_ (tetramer)) followed by motif type (P (parallel) or A (antiparallel) or H (hybrid)) which is finally followed by a number (assigned serially based on first come first serve basis) representing different motifs. Note that in the case of discontinuous folds, the fourth alphabet is indicated as "X" since it is not feasible to identify the strand orientation due to the discontinuous nature of the motif. For instance, a continuous monomeric motif with parallel strand orientation is indicated as QCMP followed by a number (**Figure 3A**). However, a discontinuous monomeric motif is represented as QDMX followed by a number (**Figure 4A**).

### Continuous motifs

Among the 33 continuous quadruplex motifs, the antiparallel motif (15) is dominantly seen, followed by hybrid (12) and parallel (6) motifs. There are two parallel (QCMP), six antiparallel (QCMA), and six hybrid (QCMH) motifs seen under the monomeric continuous (QCM) category. One can visualize from **Figure 3** that the type and the direction of the connecting loops (lateral/propeller/diagonal) bring diversity to these motifs. For instance, although QCMP1 and QCMP2 motifs are connected through a propeller, they are different due to the direction of the propeller loop. While the QCMP1 motif is connected by clockwise propeller loops, QCMP2 is connected by counterclockwise propeller loops; thus, they become mirror images of each other in terms of the loop (**Supplementary Figure S1A**). Note that the G-core maintains the same helicity in both the cases. Similarly, QCMA1 and QCMA2 (**Figure S1B**), as well as QCMA3 and QCMA4 (Figure S1C), are mirror images. QCMA1 and QCMA3 are examples of how the type of loop leads to a different motif. Although both of them have two lateral loops, the presence of a third lateral loop in QCMA1 instead of a diagonal loop in QCMA3 leads to a different monomeric antiparallel quadruplex motif. In a similar fashion, the dimeric, trimeric, and tetrameric motifs can be defined as shown in **Figure 3B-D**. Among the described multimeric motifs, QCDA2 and QCDA3, QCDH1 and QCDH2, QCT_r_P1 and QCT_r_P2, and QCT_r_H1 and QCT_r_H2 are the mirror image motif pairs. One can envisage that the remaining motifs can also have their mirror image based on the sequence and environment. One more well-known fact is that the parallel motifs are always connected by propeller loops, and the tetrameric motifs don’t have any propeller or diagonal or lateral loops.

### Discontinuous motifs

There are 15 discontinuous quadruplex motifs known as of now. Since one to three adjacent quartet residues are not connected, it is not possible to classify them based on the strand orientation into parallel, antiparallel, or hybrid, unlike continuous motifs. However, based on the number of strands, they can be classified into monomer, dimer, and trimer. There are 6, 3, 6 monomeric, dimeric, and trimeric discontinuous motifs observed so far. Intriguingly, discontinuous quadruplex motifs can be derived from continuous quadruplex motifs. One such example is QCMP1 and QDMX4, wherein the removal of connectivity between the last two residues of QCMP1 leads to QDMX4. Thus, theoretically, each continuous motif can lead to one or more discontinuous motifs. Similar to QDT_r_X4 and QDT_r_X5, which are mirror image motifs, other discontinuous motifs can also have their own mirror image motifs.

The different quadruplex motifs presented here may have different influence on interaction with the ligand molecules. Indeed, it has been discussed in a recent study that 8 distinct G-quartets are possible based on the combination of anti/syn glycosyl conformations which impacts the quadruplex groove widths, thereby affecting the ligand binding [69]. Since the complexity further increases when G-quartets are stacked on top of each other due to the inclusion of different types of loops, resulting in different motifs which may further influence the ligand binding with the quadruplex.

### Modular representation of DNA quadruplex folds

#### i. *Example illustrating the construction of simple quadruplex fold from the motif*

**Figure 5** shows a detailed representation of how the continuous parallel, antiparallel, and hybrid quadruplex motifs (**Figure 3**) can be used as a scaffold to construct simple quadruplex folds. For instance, QCMP1 is used for the modular construction of a monomeric parallel quadruplex fold (**Figure 5A (Left**)) by simply adding a G-quartet without affecting the overall quadruplex motif structure. Since QCMP1 has only G-quartets, the third quartet can be added either on the top, bottom, or middle to form a fold. In the same way, more G-quartets can be added to increase the length of the quadruplex fold. However, when a non-G-quartet is present, it has to be placed in the appropriate place, unlike in the case of all G-quartets. In a similar fashion, the examples given in **Figure 5A (middle)** and **5A (right)** illustrate the modular construction of monomeric antiparallel and monomeric hybrid folds using QCMA6 and QCMH4, respectively. One can also construct the dimeric, trimeric, and tetrameric continuous quadruplex folds using the appropriate motifs. The examples given in **Figure 5B (Left)**, **Figure 5B (Middle),** and **Figure 5B (Right)** represent the construction of the dimeric fold using parallel **QCDP1**, antiparallel **QCDA4,** and hybrid **QCDH1** motifs, respectively. **Figures 5C** and **5D** are examples of trimeric and tetrameric fold construction, respectively. A total of 12 different examples of deriving a simple quadruplex fold from the motifs are illustrated in **Figure 5**.

**Figure 5.**
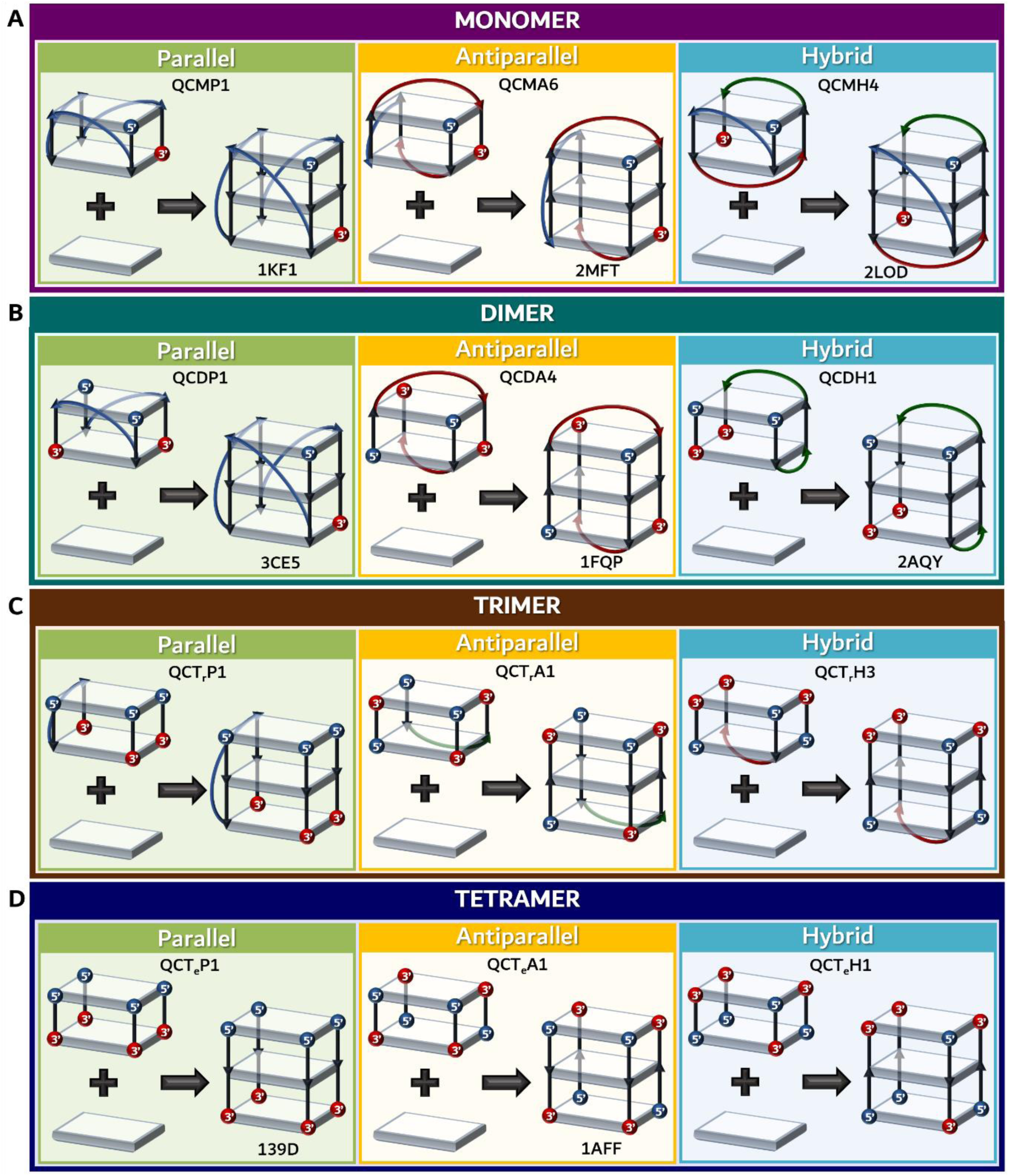
Examples illustrating the modular construction of A) monomeric, B) dimeric, C) trimeric, and D) tetrameric (Left) parallel, (middle) antiparallel, and (Right) hybrid quadruplex folds using the continuous quadruplex motifs described in **Figure 3**. Note that the name of the continuous monomeric motif is given alongside each panel. A representative PDB ID is given adjacent to each fold, wherever applicable.

#### ii. *Example illustrating the construction of complex quadruplex fold from the motif*

Unlike the above cases, the construction of a complex quadruplex fold is not straightforward. Such a situation arises when at least one of the motifs is discontinuous. In order to construct a discontinuous quadruplex fold, the quartets common to two motifs that form the fold should be superimposed to get the complete fold. Unlike in the continuous fold, wherein the G-quartet can be added in the middle or top, or bottom, here, two quartets should be overlayed to maintain the directionality of the fold. Further, based on the nature of the quadruplex fold, one or more additional loops are required to complete the fold which is dictated by the sequence. For instance, in **Figure 6A**, the quadruplex fold can be established by a continuous motif QCT_r_P1 and discontinuous motif QDT_r_X2, wherein these two motifs are superimposed with respect to the common quartet (indicated in gray colored stripes in **Figure 6A** and **6B**). Note the two new propeller loops in the final fold (colored blue) that make the fold monomeric. Interestingly, the addition of QCT_r_P1 and QDT_r_X2 motifs without the formation of 2 propeller loops makes a trimeric fold. It is noteworthy that placing the QDT_r_X2 motif on the top of QCT_r_P1, which is the reverse of the above, would result in a different fold (no such quadruplex fold structures are observed in PDB as of now) depending on the sequence and environmental condition. Yet another such example is shown in **Figure 6B**. An example of creating a quadruplex fold with more than two quadruplex motifs is shown in **Figure 6C** (**See Figure 6C for details**).

**Figure 6.**
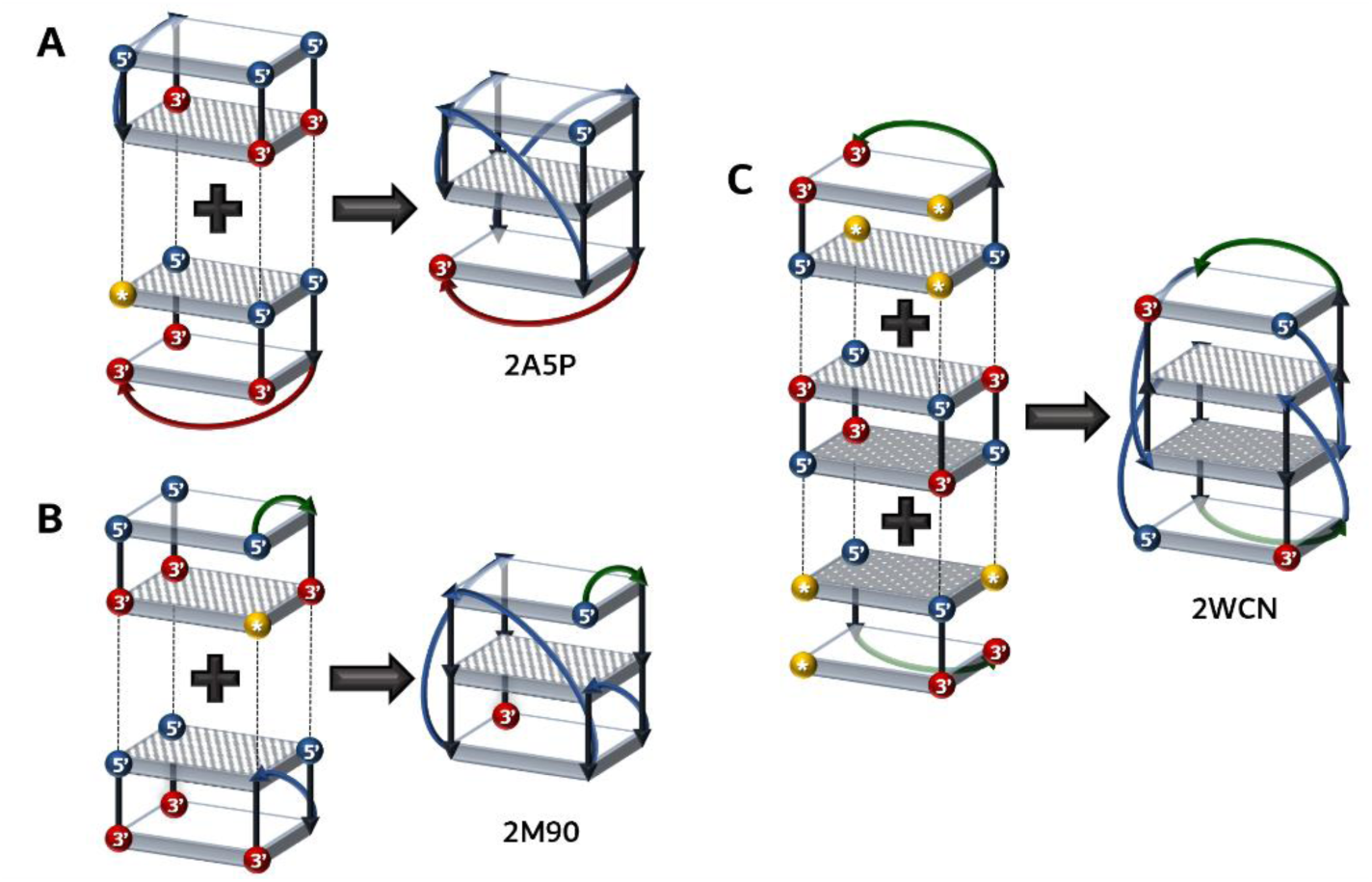
Examples illustrating the modular construction of monomeric (A and B) and dimeric discontinuous (C) quadruplex folds using the quadruplex motifs described in **Figures 3 and 4**. A) and B) Constructing a quadruplex fold with the use of one discontinuous and one continuous motif. Note the formation of two different folds by just changing one of the motifs, *viz.*, while the continuous motif is the same between the 2 cases, the discontinuous motif is different. C) An example of forming a higher-order fold is shown by considering three motifs (one continuous motif amid two identical discontinuous motifs). Note the formation of four new propeller loops (colored blue) to complete the fold. (A-C) Dotted lines indicate the point of linkage between 2 different quadruplex motifs. The overlapping G-quadrates are indicated by striped (A-C) and checked (C) rectangular slabs. A representative PDB ID is given adjacent to each fold.

#### iii. *Example illustrating the construction of higher-order quadruplex structures using the motifs*

Higher-order quadruplex folds are more realistic in a biological context [42, 70, 71], and there are 13 such quadruplex topologies seen in PDB (labeled as "Higher-order" in **Supplementary Table S1**). The analyses of the known quadruplex fold structures indicate two types of higher-order topologies: homo and hetero. In the homo quadruplex folds, more than one repetition of a continuous or discontinuous motif occurs. However, the hetero quadruplex folds are formed by the combination of more than one continuous and/or discontinuous motif. In the higher-order quadruplex, two or more same or different quadruplex motifs are connected through linker (l) loop(s), which can be considered a special case of a bulge loop that connects two motifs. One such example is given in **Figure 7A**, wherein two higher-order homo quadruplex folds are formed by two units of the same QCMP1 motif, which is dictated simply by the difference in the orientations between the two motifs in the Y- and Z-directions. While there are two thymines (T) in the linker loop for the fold given in **Figure 7A (top)**, there is either a single T or two "T" s or two adenines (A’s) in the linker loop for the fold given in **Figure 7A (bottom)**. This suggests the negligible role of the loop residues in dictating the formation of these folds. This also suggests that the relative orientation between two motifs (homo or hetero) may lead to many combinations of higher-order quadruplex folds depending on the rotation along the X- and/or - Y and/or Z axes. When the number of motifs increases, it invites more complications in the higher-order arrangement of the quadruplex topology. In addition, the position of linker loop nucleotides in the sequence may lead to different arrangements between the motifs. An example of having more than two continuous motifs forming a hetero quadruplex topology is given in **Figure 7B**, wherein the end motifs of the quadruplex fold are continuous dimer parallel, whereas the middle one is a tetramer antiparallel continuous motif. One can expect more complexities with many such combinations of parallel, antiparallel, and hybrid motifs. Homo higher-order quadruplex fold can be formed with two discontinuous motifs **(Figure 7C)**. A hetero quadruplex fold can also be formed with discontinuous and continuous motifs, as given in **Figure 7D**.

**Figure 7.**
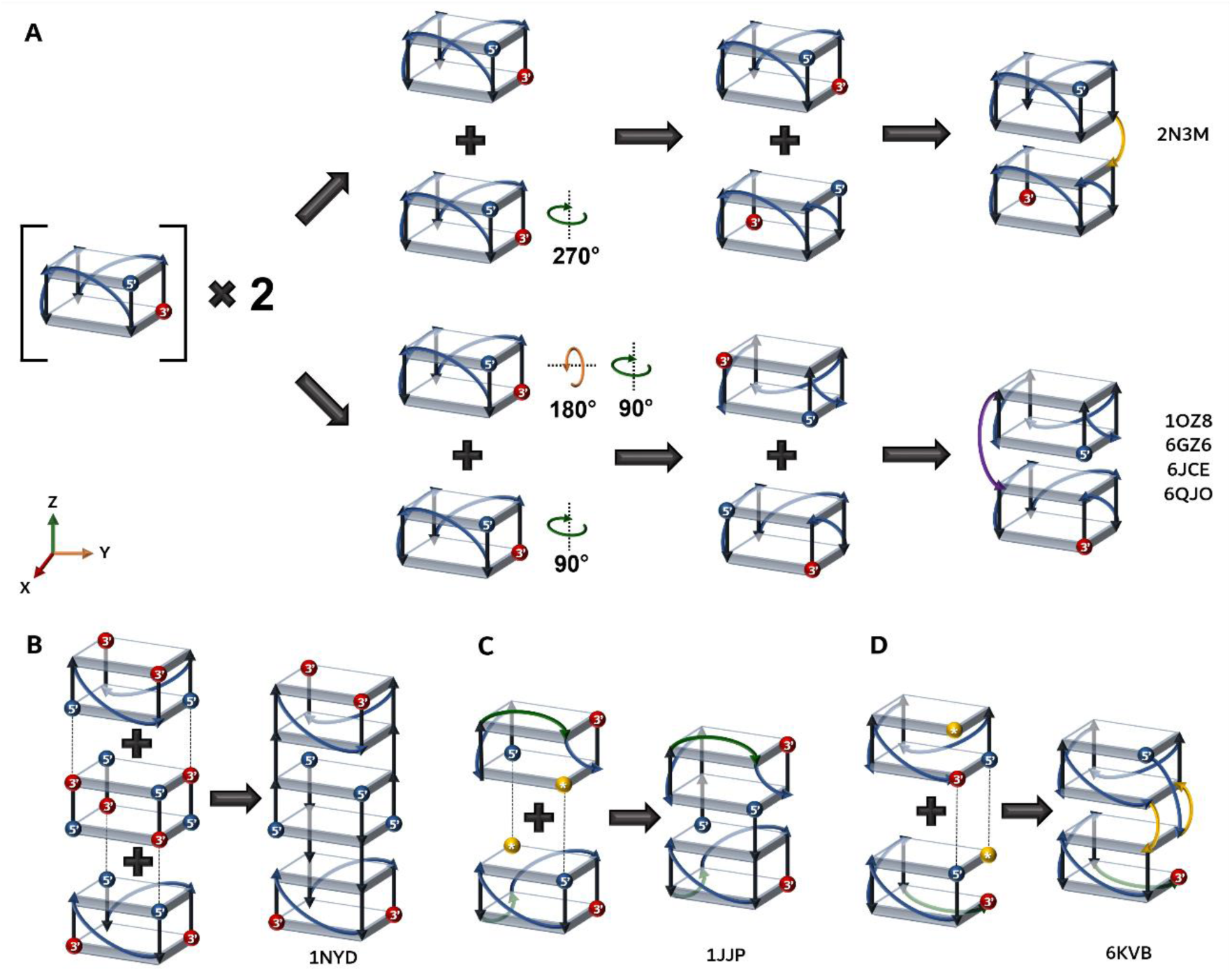
Schematic illustration of constructing a higher-order (A and C) homo and (B and D) hetero quadruplex folds. A) Two units of QCMP1 forming two different higher-order stacked quadruplex folds through a linker loop are shown. Note the possibility of having two different higher-order folds out of the same scaffold motif by just changing their relative orientations governed by the rotation around X, Y, and Z axes (indicated in circular arrows; X-, Y- and Z-axes arrows are colored red, yellow and green respectively). Formation of a B) hetero higher-order quadruplex fold using two different continuous motifs (QCDP1 and QCT_e_A2), C) homo higher-order quadruplex using two same discontinuous motifs (QDMX6), and D) hetero higher-order quadruplex using two discontinuous motifs (QDMX4 and QDMX5). A representative PDB ID is given adjacent to each fold.

### Devising a universal nomenclature to define the folding architecture of quadruplexes

A total of 49 unique DNA quadruplex folds (**Figures 8-10**) are identified from the analysis of 388 PDBs. Among the 49 folds, 23, 13, and 13 are continuous, discontinuous, and higher-order folds. Although these folds are predominantly formed by G-quartets (**Figure 1A**), non-G-quartets like T-T-T-T (PDB ID: 7D31) [72, 73], C-C-C-C (PDB ID: 6A85) [73], A-T-A-T (PDB ID: 5LS8) [74], G-C-G-C (PDB ID: 7CV4) [75], C-A-G-A (PDB ID: 6ZX7) [76], and G-G-A-T (PDB ID: 5VHE) [77] are also found. Out of the 223 sequences considered for the analysis, the number of sequences preferring continuous parallel folds (three folds) is more compared to the antiparallel and hybrid folds, *viz.*, 81 unique sequences among 143 structures prefer parallel fold. Thus, it is clear that different DNA sequences assume an identical fold. Interestingly, an identical sequence taking different folds is also observed, indicating the role of environmental conditions in dictating the quadruplex fold. For instance, the sequence "GGGTTAGGGTTAGGGTTAGGG" assumes both parallel (PDB ID: 1KF1) [78] and antiparallel (PDB ID: 143D) [79] folds. When this quadruplex forming sequence has overhangs, it further assumes three more folds (PDB IDs: 2GKU [80], 2JPZ [81], 2MBJ [82]).

**Figure 8.**
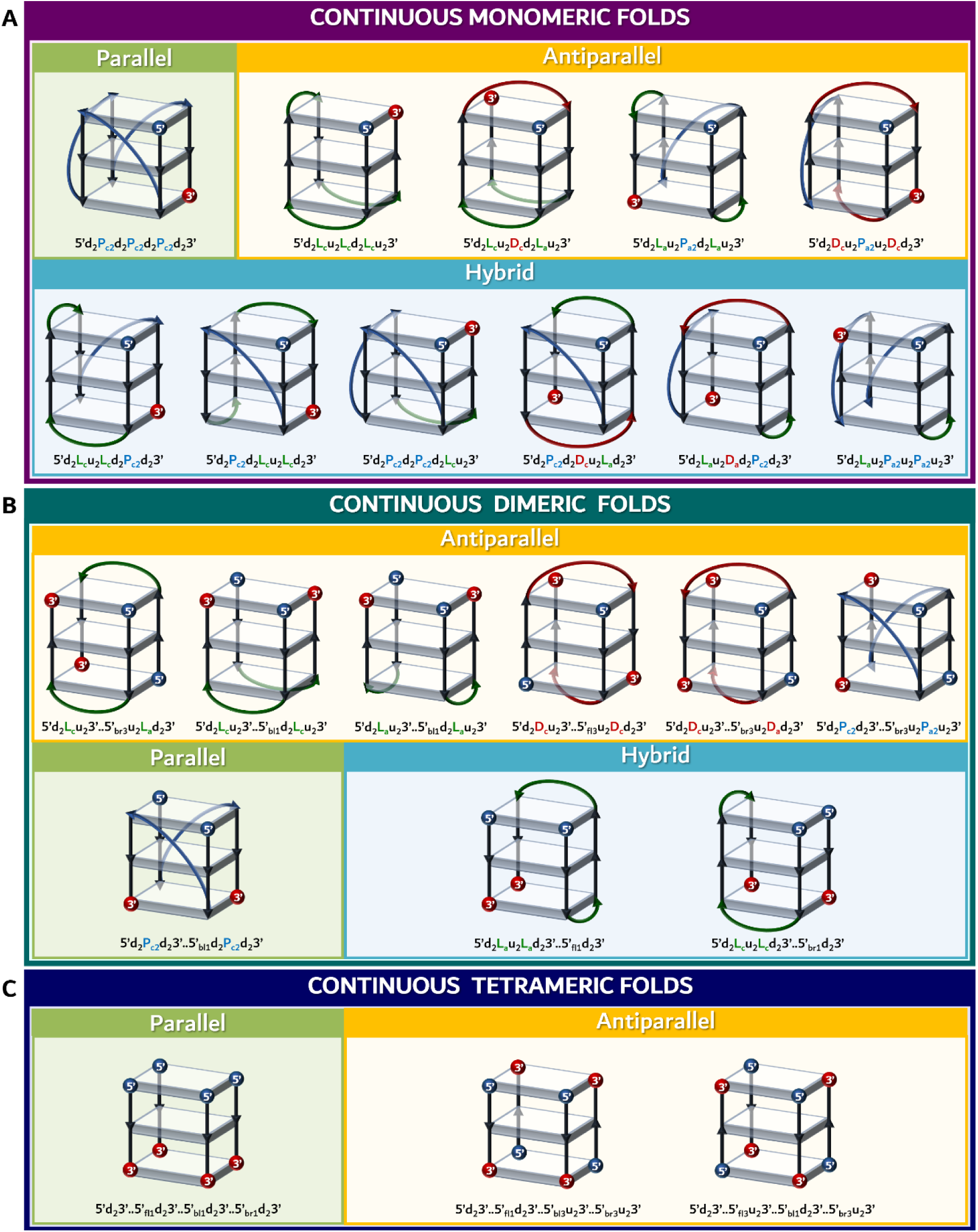
Schematic representation of unique continuous DNA quadruplex folds. For illustration purposes, each fold is represented with only three quartets; Refer to Supplementary **Table S1** for the details about the PDB IDs, sequence, number of quartets, *etc.*, corresponding to each fold described. Note that the corresponding alphanumeric nomenclature is indicated adjacent to each fold.

An alphanumeric nomenclature has been devised here to represent the quadruplex folds, as discussed below. The nomenclature starts with the 5’ end of the quadruplex strand. By default, 5’ indicates the first (_1_) quartet present on the right-hand side (_r_) front face (_f_) of the quadruplex fold. Following this definition, the suffix ’_f_’ or ’_b_’ is added to the 5’ to indicate whether the 5’ end starts from the front or back side of the quadruplex fold, respectively. This is then followed by a suffix ’_l_’ or ’_r_’ to indicate the location of the 5’end in the left- or right-hand side of the quadruplex, respectively. Finally, a numerical value is added in the suffix to represent the position of the quartet (with the numbering scheme going in ascending order from the top of the quadruplex fold). For instance, 5’_bl3_ refers that the 5’ end of the quadruplex strand starts from the third quartet of the back face located on the left-hand side of the quadruplex. Following this, the chain direction is indicated by a diagonal loop (D_c/a_) or lateral loop (L_c/a_) or bulge loop (B) or propeller loop (P_(c/a)n_), or the number of quartet residues the chain has to travel down (d_n_) or the number of quartet residues the chain has to travel up (u_n_). While the suffix ’_c/a_’ in D, L, and P refers to the clockwise or anticlockwise direction of the chain, ’_n_’ in P, d, and u indicates the number of quartet residues. Finally, the naming scheme ends with 3’ to denote the 3’ end of the quadruplex strand. This nomenclature can readily be used for referring to all the continuous (**Figure 8**) and discontinuous (**Figure 9**) quadruplex folds. For instance, 5’d_2_D_c_u_2_P_a2_u_2_D_c_d_2_3’ refers to the monomeric continuous fold given in the right-most end of the first row of **Figure 8A**. In a similar fashion, each strand of the intermolecular quadruplex can be referred. Here, the different strands of the quadruplex fold are differentiated by ’..’. The same nomenclature can be extended for the higher-order quadruplex folds, wherein the intertwined quadruplex folds pose more challenges. Due to such complexity, the alphanumeric nomenclature is not mentioned here for the higher-order quadruplexes (**Figure 10**).

**Figure 9.**
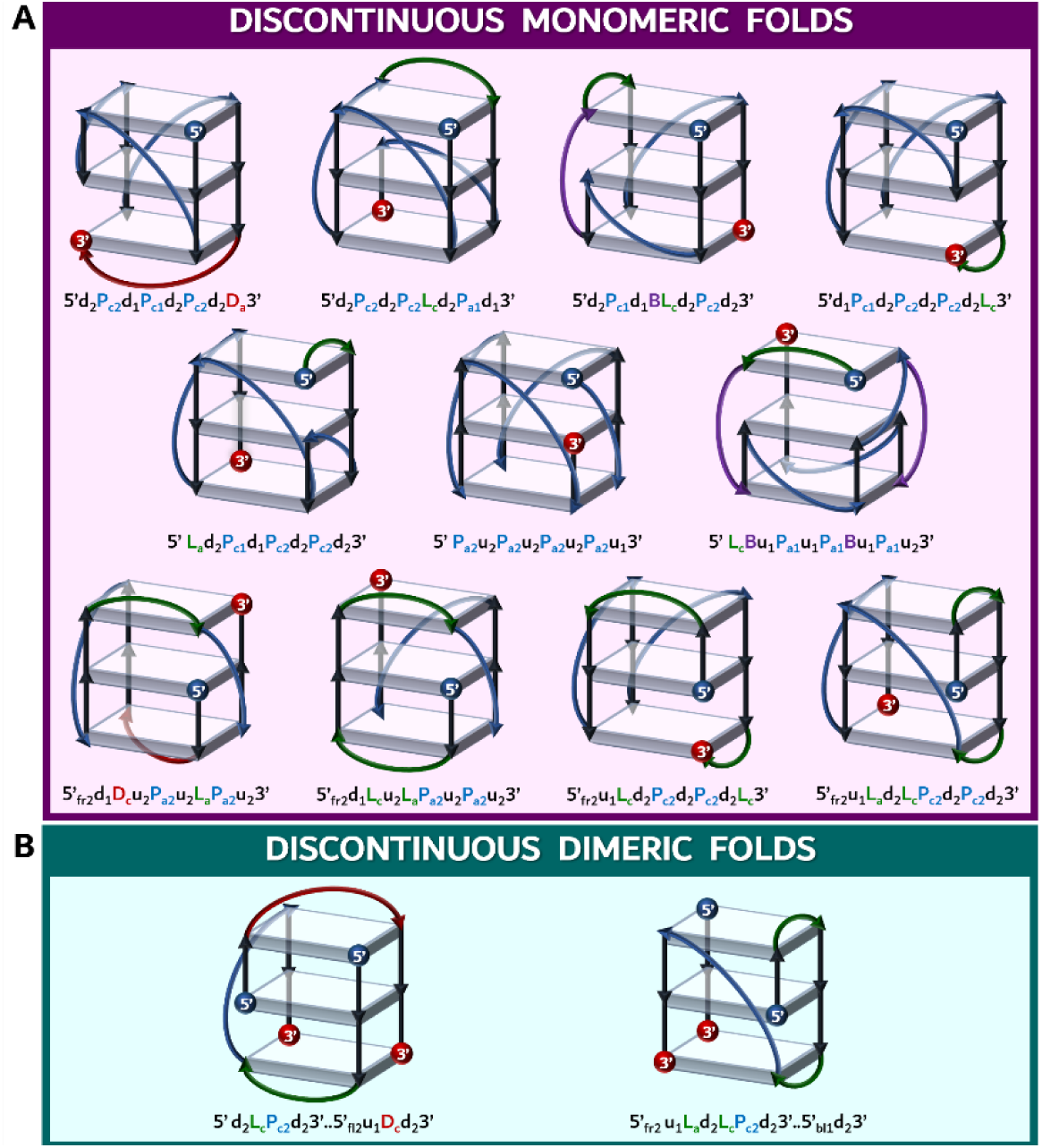
Schematic representation of unique discontinuous DNA quadruplex folds. For illustration purposes, each fold is represented with only three quartets; Refer to Supplementary **Table S1** for the details about the PDB IDs, sequence, number of quartets, *etc.*, corresponding to each fold described. Note that the corresponding alphanumeric nomenclature is indicated adjacent to each fold.

**Figure 10.**
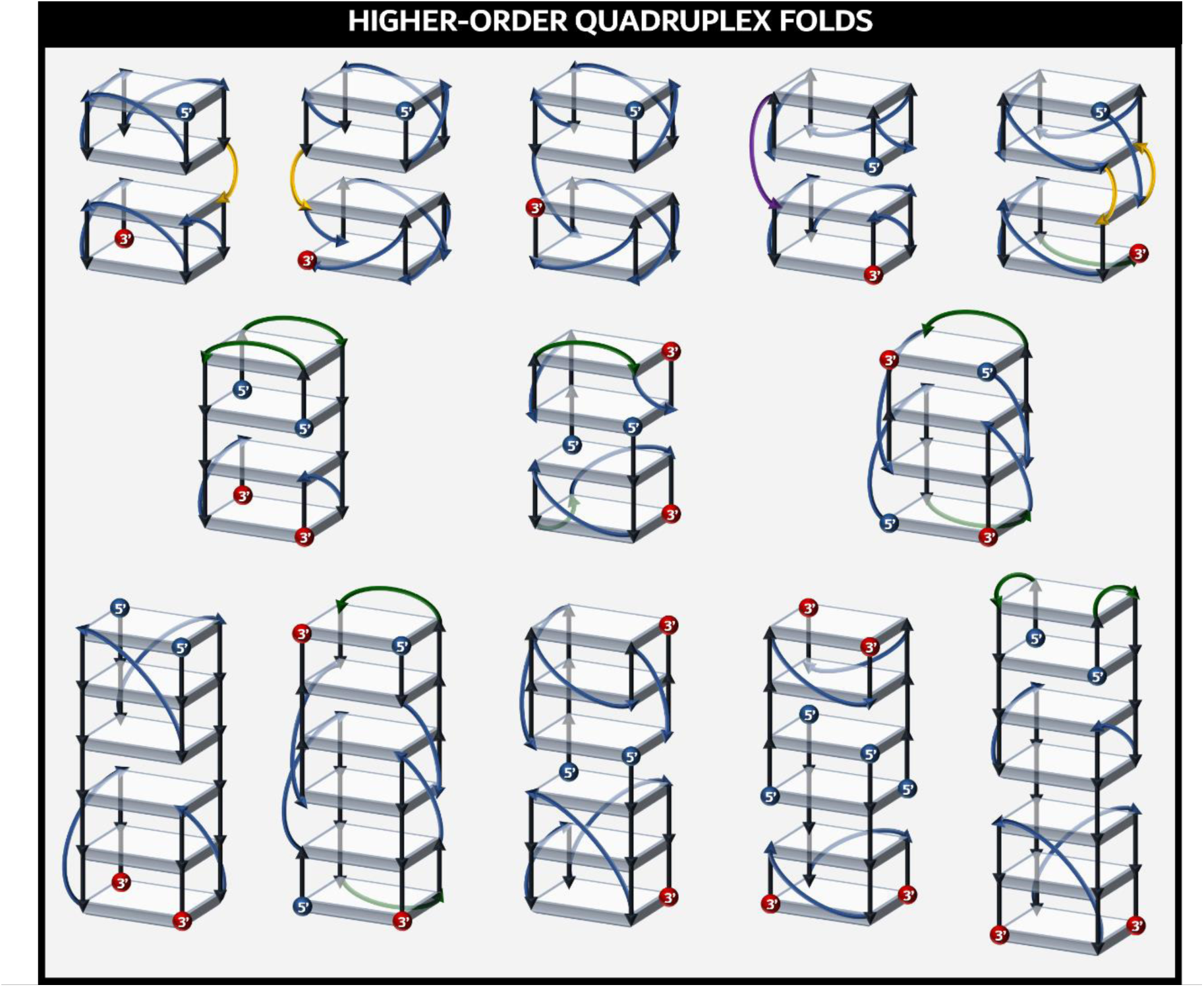
Schematic representation of unique higher-order DNA quadruplex folds. Refer to Supplementary **Table S1** for the details about the PDB IDs, sequence, number of quartets, *etc.*, corresponding to each fold described.

Among the 49 DNA folds discussed here, 8 of them are found to be common with RNA quadruplex folds [27]. However, some folds are specific for RNA quadruplex, which are not discussed here.

### Conformational angle preference for different quadruplex folds

The conformational angle preference of the quadruplex folds is analyzed individually for G-quartet forming residues and the loop residues. The conformational angles corresponding to each PDB ID (given in **Supplementary Table S1**) deposited in NDB [61] are downloaded and compiled in an Excel file. These conformational angles are then used to generate the density plots for the torsion angles with the help of Gnuplot [67]. The backbone conformational angles α (O3’-P-O5’-C5’), β (P-O5’-C5’-C4’), γ (O5’-C5’-C4’-C3’), δ (C5’-C4’-C3’-O3’), ε (C4’-C3’-O3’-P) and ζ (C3’-O3’-P-O5’) and, the glycosyl conformation angle χ (C4-N9-C1’-O4’) (**Supplementary Figure S2A**) of the quartet guanine residues exhibit preference for *gauche-*, *trans*, *gauche+*, *trans*, *trans*, *trans-* and *anti* (**Figure S2B-G**) respectively with a few exceptions. The *trans* conformation for ’δ’ indicates the C2’-endo sugar puckering preference, and only a minor population of C3’-endo (*gauche+*) is seen. This is in contrast to RNA-quadruplex, wherein both C2’-endo and C3-endo sugar puckers are found to be equally favorable [27]. Further, χ shows a preference for *anti*-conformation in the parallel G-quartet (**Figure S2H**) as it naturally facilitates the hydrogen bonding patterns between the Gs. However, antiparallel and hybrid quadruplexes show a preference for *+syn* conformation in order to maintain the hydrogen bonding pattern (**Figure S2H**). While 2 of the guanines take *+syn* conformation in the antiparallel quadruplex, one of the four guanines in the hybrid quadruplex takes *+syn* conformation.

The loop residue conformational angles α, β, γ, ζ and χ predominantly prefer *gauche-*, *trans*, *gauche+*, *gauche-*, and *anti* conformations, respectively (**Figure S3**). Interestingly, ε takes a wide range of conformational angles in the range of 180-300° (**Figure S3E**). A significant population of *gauche+*, *trans*, and *gauche+*/*trans* is also seen for α, γ, and ζ respectively, similar to the quartet guanine residues (**Figure S2-S3**). δ value further confirms the C2’-endo sugar pucker (**Figure S3**).

## Conclusions

A detailed analysis of experimentally derived quadruplex structures is carried out here to understand the architecture of DNA quadruplexes. The modular nature of quadruplex architecture is established here by defining quadruplex motifs. Utilization of these motifs as scaffolds for simple as well as higher-order quadruplex folds is also demonstrated. Such knowledge about the quadruplex topologies will be helpful in the programmed design of quadruplex folds, motif-specific ligand design, and understanding the interaction with the proteins. One can further explore the sequence specificity of the quadruplex motifs and utilize them in the genome-wide prediction of quadruplex motifs with the help of machine learning approaches. Further, a universal nomenclature has been devised here to represent the quadruplex folds. Such an alphanumeric representation of quadruplex folds may be helpful in training machine learning models to predict and model the quadruplexes.

## Acknowledgments

The authors thank Sathyaseelan for suggestions on PDB data collection. The authors thank the Indian Institute of Technology Hyderabad for their computation resources. SS and PPU thank MoE and CSIR, respectively, for the fellowship.

## Author contributions

SS collected the PDB data and carried out quadruplex motif and fold analyses. PPU collected the PDB data, participated in fold analysis, and carried out conformational angle analyses. PK carried out fold and loop residue analyses during the early stage of the project. SS and TR wrote the manuscript. TR designed and supervised the project.

